# Immunities Specific to Both of the M Protein Ectodomain and RBD Synergize to Confer Cross-protection against SARS-CoV-2 Infections

**DOI:** 10.1101/2023.07.31.551223

**Authors:** Yibo Tang, Kaiming Tang, Yunqi Hu, Zi-Wei Ye, Wanyu Luo, Cuiting Luo, Hehe Cao, Ran Wang, Dejian Liu, Cuicui Liu, Xingyi Ge, Yaoqing Chen, Shuofeng Yuan, Lei Deng

## Abstract

The effectiveness of the prototypic SARS-CoV-2 vaccine largely decreased overtime against the emerging virus strains, necessitating the universal vaccine development. The most abundant structural membrane (M) protein is highly conserved in amino acid sequence, which arouses our research interests in developing a universal immunogen based on it. Serological analysis showed that IgG responses specific to its N-terminal peptides can be strongly detected in many serum samples from both convalescent patients and vaccinees receiving inactivated vaccines, indicating the potential existence of human B-cell epitopes in reactive peptides. Microneutralization assays showed that the N-terminal peptide S2M2-30-specific hyperimmune serum was capable of cross-neutralizing the authentic viruses including wild-type HKU-001a, B.1.617.2/Delta, and Omicron subvariant BQ.1.1, and synergized with RBD-specific serum in reinforcing antiviral activities. Strong S2M2-30-specific immunities elicited in hACE2-transgenic mice could effectively inhibit B.1.1.7/Alpha (UK) infections. Our results suggest the potentiality of conserved M peptides as vaccine targets for conferring cross-protections against sarbecoviruses.

## Introduction

Currently used SARS-CoV-2 vaccines have been developed based on targeting the S protein or its RBD subdomain from the prototypic strain for inducing neutralizing immuno-protection in humans ^1^. SARS-CoV-2 viruses constantly change through mutation, occasionally resulting in new variants that allow viral escape from herd immunity and spread more easily. Increasing investigations revealed that the specific immune responses elicited by shots with these marketed vaccines conferred dramatically reduced neutralizing protection against the more contagious Omicron variant strains and the recently emerging XBB and BQ.1 variant strains that are already rising above others in spread ^2–6^. The sobering facts that SARS-CoV-2 viruses most likely continue to circulate indefinitely among the worldwide population suggest the urgent need for a universal vaccine that can induce lasting broadly protective immunity.

M protein is the most abundant structural protein in SARS-CoV-2 virion particles and plays versatile and pivotal roles in directing virion assembly, morphogenesis, and antagonizing MAVS-mediated interferon responses ^7–9^. Coronavirus M protein ectodomain has ever been verified to participate in the viral attachment to receptors, facilitating viral entry into the host cells ^10^. N-terminal ectodomain-specific IgG responses were detected in human convalescent serum samples, indicating the presence of human B-cell epitopes in the N-terminal ectodomain ^11^. The sufficient space between neighboring S trimers on the SARS-CoV-2 virion surface allows the M protein ectodomain-specific antibodies to penetrate and bind to M proteins beneath S proteins on the virion surface. The bindings of these antibodies to virion would conduce to the mitigation of the viral attachment to host cells ^12, 13^, presumably, could also potentially sterically impede the post-fusion state change of S proteins, then the viral neutralizations could ensue. This hypothesis was made in the light of a phenomenon that once bound to the influenza hemagglutinin the antibody CR9114 would also somehow switch off the viral neuraminidase activities ^14, 15^. In the infected cells, plenty of M proteins that miss the incorporation into virions are destined for anchoring on the cellular plasma membrane and thereafter are bound to be the targets for specific antibody-mediated cellular clearance. Of note, M protein ectodomain from sarbecovirus subgenus strains are highly conserved in the amino acid sequence and hold potentiality for inducing cross-protection against both SARS-CoV-1 and SARS-CoV-2. These inferences above elicited our interest in the immunological investigations of the M protein ectodomain-based immunogens.

Inactivated SARS-CoV-2 vaccines containing M proteins for sure have been given over a billion doses worldwide ^16^. We are curious about whether inactivated vaccines would induce strong M protein ectodomain-specific antibody responses and especially the protective effects thereof. For now, as never has a vaccine immunogen solely based on the ectodomain from SARS-CoV-2 M protein been reported, we have no clue yet if its specific immunity is protective or not. Whether its specific antibody response contributes to COVID-19 immunopathology via antibody-dependent enhancement (ADE) is also a matter of great concern. Non-neutralizing or sub-neutralizing antibodies can cause enhanced viral infections of monocytes via FcγRIIa-mediated endocytosis. Besides, antibodies complexed with viral antigens inside airway tissues, the ensuing pro-inflammation cytokine responses, and immune cell recruitments present another main ADE mechanism ^17^.

An essential precondition for the peptide-based vaccine to induce high-affinity virus-specific antibody responses is that the incorporated peptides in the immunogen can reproduce the native conformation as presented in the virion. The structure of the extreme N-terminal peptide of M protein in solution is technically difficult to be resolved and its exact amino acid sequence length is also uncertain ^9^. It was reported that M proteins form homo-dimer and higher-order oligomers in virions ^9^, raising a scientific question if the induction of protective antibodies is quaternary structure-dependent. Only in the case that the N-terminal ectodomain in virion particle presents as a flexible loop structure, can the design of an effective N-terminal peptide-based vaccine not be strictly constrained by conformation requirement. In this study, N-terminal peptides-specific IgG antibody responses can be positively detected in most of the human convalescent serum and the serum samples from vaccinees who ever received three shots with CoronaVac COVID-19 inactivated vaccines. We found a peptide, named as S2M2-30, showed strong immunogenicity. The immunizations of human angiotensin-converting enzyme 2 (hACE2)-transgenic mice with the S2M2-30-based immunogens demonstrated the effective cross-protectivity of S2M2-30-specific immunity. Reference could be made to the adaptive immunology outcomes from this study for comprehensive evaluation of the authentic functions of the M protein-specific herd immunities.

## Results

### Designs of N-terminal peptides from coronavirus M protein

Unraveling the exact ectodomain of M protein facilitates our designs of immunogens for inducing protective antibody responses, the circumstantial evidence obtained from the solved SARS-CoV-2 M protein structures suggests the amino acid sequences 2-19 and 72-79 (Figure 1 A). As the structure of the extreme N-terminal peptide has not been solved yet, we employed the popular computational tool Alphafold-2 to model the structure of single-chain M polypeptide from SARS-CoV-2, SARS-CoV-1, and MERS-CoV. This peptide from any of the indicated coronavirus was predicted as a loop structure and perhaps wiggled on the membrane surface (Figure 1 B). The consensus amino acid sequence of 2-30 or 71-80 M peptide was deduced (Figure 1 C). We found that the sarbecovirus M protein ectodomain is an evolutionarily conserved region but is totally unlike the amino acid sequence from MERS-CoVs. A few mutations like D3G/N and Q19E in Omicron SARS-CoV-2 M protein possibly contribute to stronger viral fitness in human hosts and also facilitate viral evasion from immunosurveillance by impairing paratope complementarity of high-affinity antibodies specific to this domain. Herein, an N-terminal peptide from MERS-CoV M protein was designed for the negative control use in the subsequent immunization studies.

**Figure 1.**
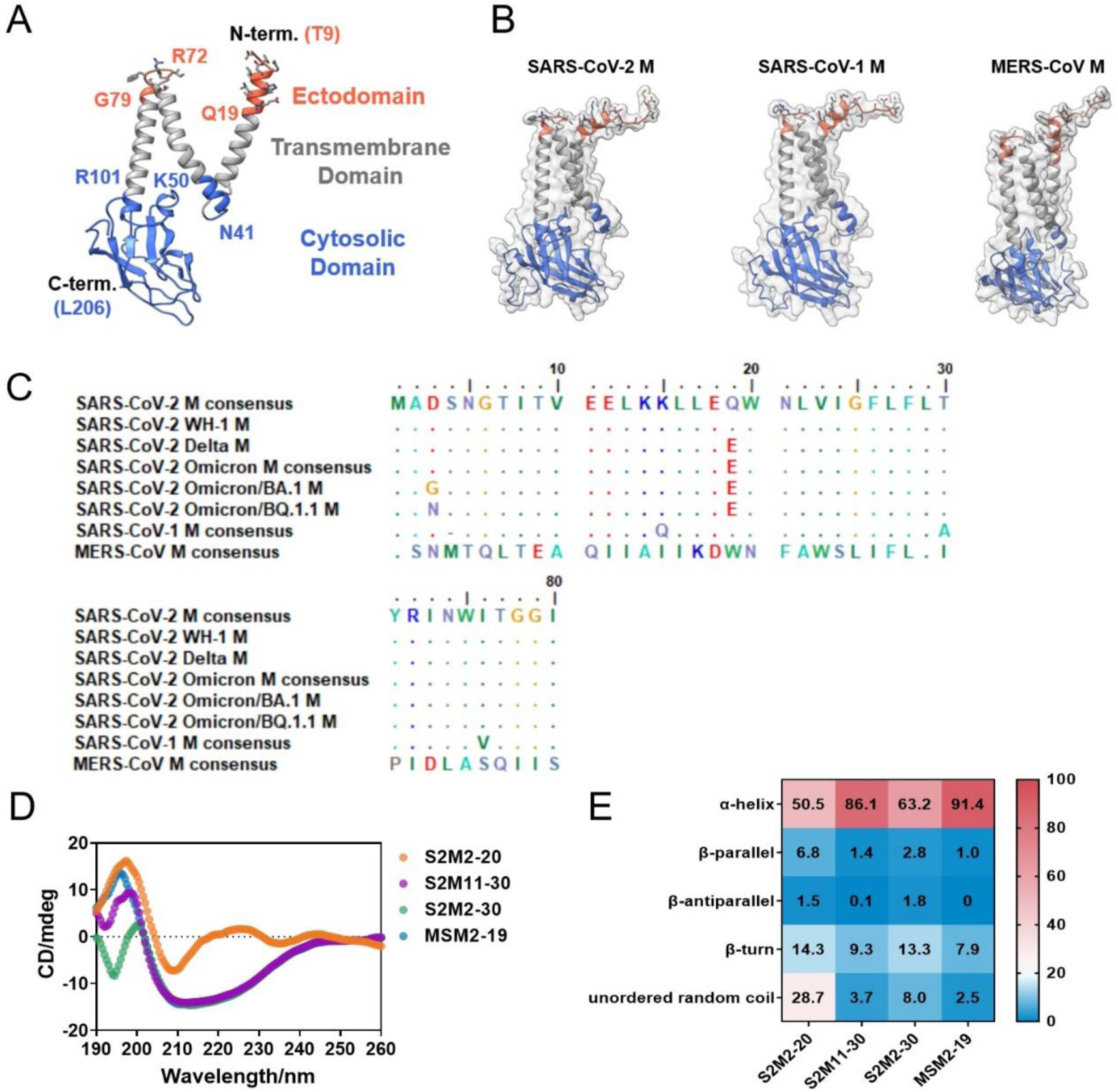
Analysis of M protein ectodomain. (A) Ribbon model of SARS-CoV-2 M protomer (PDB ID: 7VGR). (B) The structure predictions of full-length M proteins from SARS-CoV-2, SARS-CoV-1, and MERS-CoV by using AlphaFold-2. (C) The amino acid sequence alignment of M protein ectodomain from different coronaviruses. (D) The CD spectra detection of S2M2-20, S2M11-30, S2M2-30, and MSM2-19 in the solid state for secondary structure analysis. (E) The output data of CD spectra analysis indicates the proportion of each type of protein secondary structure.

The peptide S1M2-19 from SARS-CoV-1 M protein, MSM2-19 from MERS-CoV M protein, and several SARS-CoV-2 M ectodomain peptides in different amino acid lengths and in the presence or absence of a monosaccharide on N5, were synthesized for the uses in serological analysis and immunogenicity evaluation (Table 1). Circular dichroism detection showed that the majority of the amino acids in peptides S2M2-20, S2M11-30, S2M2-30, and MSM2-19 form alpha-helices (Figure 1 D, E). It is speculated that the rigid secondary structures in S2M2-30 are largely formed by the fragment spanning from the amino acid 11 to 30.

**Table 1.** Amino acid sequences of the synthetic peptides from coronavirus M protein ectodomain. The (gly) indicates the addition of a monosaccharide at N5.

| Viruses | Peptide names | Amino acid sequences |
| --- | --- | --- |
| SARS-CoV-2 | S2M2-20 | ADSNGTITVEELKKLLEQW |
|  | S2M8-24 | ITVEELKKLLEQWNLVI |
|  | S2M11-30 | EELKKLLEQWNLVIGFLFLT |
|  | S2M2-30 | ADSNGTITVEELKKLLEQWNLVIGFLFLT |
|  | S2M74-77 | NWIT |
|  | S2M2-20-Mid | ADSNGTITVEELKKLLEQWGSGSNWIT |
|  | Gly-S2M2-20-Mid | ADSN(gly)GTITVEELKKLLEQWGSGSNWIT |
|  | S2M11-20 | EELKKLLEQW |
|  | S2M21-30 | NLVIGFLFLT |
| SARS-CoV-1 | S1M2-19 | ADNGTITVEELKQLLEQW |
| MERS-CoV | MSM2-19 | SNMTQLTEAQIIAIKDW |

### SARS-CoV-2 infections or immunizations with inactivated vaccines elicited M protein ectodomain-specific IgG responses in human

To test and compare human serum IgG reactivities to the N-terminal peptides of coronavirus M proteins, we collected 10 serum samples from unvaccinated convalescent COVID-19 patients who had been confirmed in 2020 by using SARS-CoV-2 reverse transcription polymerase chain reaction (PCR), 10 serum samples from vaccinees receiving two shots with CoronaVac COVID-19 inactivated vaccines but uninfected yet, and 10 serum samples naïve to SARS-CoV-2 from children outpatients in Beijing Children’s Hospital in the middle of 2019 as the negative controls in this study. Enzyme-linked immunosorbent assay (ELISA) results showed that most of the convalescent serum samples exhibited remarkably stronger binding activities to all peptides derived from the M proteins of SARS-CoV-2 and SARS-CoV-1, compared with either the inactivated vaccine-induced serum or the pre-pandemic serum samples (Figure S1). Two immunizations with the inactivated vaccine could weakly induce the specific IgG responses to certain M peptides. However, we did not observe the stronger serum IgG reactivity to S2M2-20-Mid where a conserved peptide comprising amino acids 74 - 77 was fused to S2M2-20. These positive serological results revealed the presence of at least one human B cell epitope in the N-terminal peptides.

The MSM2-19-specific IgG responses were weakly detected in the convalescent and inactivated SARS-CoV-2 vaccine-induced serum samples (Figure S1). To our understanding, this weakly positive serological result from either group is hard to be clearly explained, presumably, is likely the unspecific background level.

In another independent serological analysis, we incorporated an additional group comprising 20 serum samples collected from vaccinees receiving three consecutive immunizations with CoronaVac COVID-19 inactivated vaccines, and also expanded the number of serum samples in each group. Similar to previous results, the human seroconversion rate to the M protein ectodomain peptides consistently showed a high level in the SARS-CoV-2 infected population and reached 80% (Figure 2 A). We also found that the second booster immunization with the inactivated vaccines remarkably increased the seroconversion rates to M protein ectodomain peptides, as showed by its ascending from 8.3% to 80% against S2M2-30 peptide and from 39.6% to 95% against S1M2-19 peptide (Figure 2 A). As expected, the convalescent serum and the inactivated vaccine-induced serum showed strong cross-reactivities to the SARS-CoV-1 M peptide S1M2-19.

**Figure 2.**
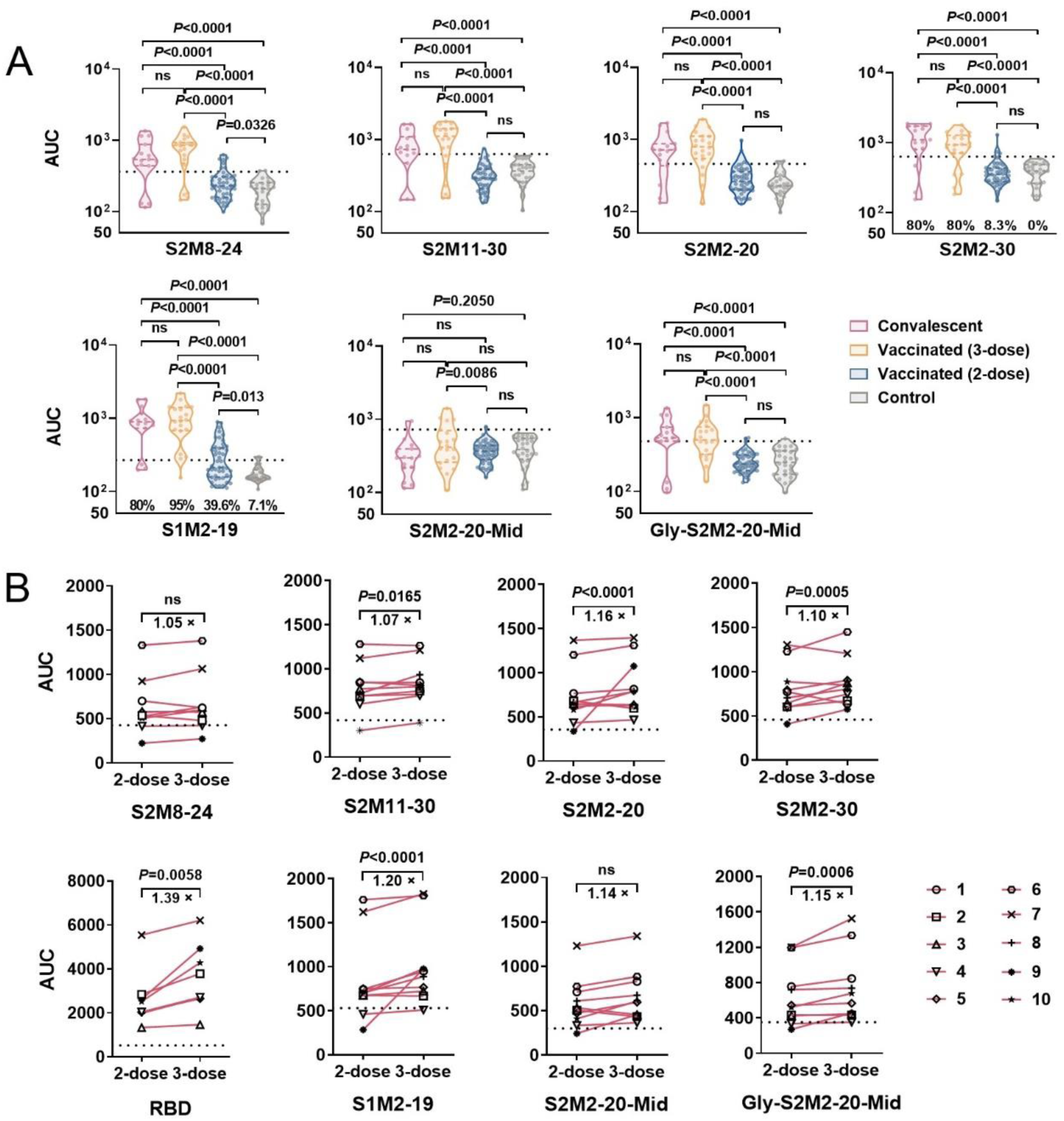
Serological evidence of M protein ectodomain-specific IgG responses in humans. (A) The area under the curve values (AUC) of specific serum antibody titers. Each point from the groups represents the specific antibody titer of an individual. Tested serum samples were collected from the COVID-19 convalescent patients (n = 15), vaccinees who ever received 2-dose (n=48) or 3-dose (n=20) of inactivated SARS-CoV-2 vaccine, and the child outpatients in the middle of 2019 as the negative controls (n = 26). Statistical significance between compared groups was assessed with an unpaired t-test. (B) Longitudinal serological analysis of serum samples from 10 vaccinees. Statistical significance was determined with two-way ANOVA. P values in all figures less than 0.05 are considered statistically significant. The dashed lines indicate the background levels that represent mean + 2 × standard derivation (SD) of the AUC values of control groups.

In order to exactly evaluate the changes in M peptide-specific serum IgG titers from the same vaccinee individuals between two follow-up periods, after the second and third immunization with inactivated vaccines, respectively. We found that the measured IgG titers were on the rise after the third immunization (Figure 2 B). The serological investigations above indicate that M protein ectodomain peptides are immunogenic in humans, and their specific IgG responses have already been prevalent among populations who were infected with the SARS-CoV-2 virus or ever received three immunizations with inactivated vaccines.

### Hyperimmune serum with both specificities to the M peptide and RBD protein exerted synergistic *in vitro* antiviral effects

To evaluate the immunogenicity of the M protein ectodomain peptides in conjugation with a protein carrier, the BALB/c mice were intraperitoneally immunized three times with peptide-keyhole limpet hemocyanin (KLH) conjugates emulsified with incomplete Freund’s adjuvant (Figure S2 A). The SARS-CoV-2 M peptides that strongly reacted with convalescent serum IgG showed varying capacities of inducing IgG responses in BALB/c mice (Figure 3 A, B). Immunizations with S2M2-30-based conjugates exhibited the highest efficacy in inducing M peptide-specific IgG responses. Whereas, the shorter M peptides from SARS-CoV-1 and SARS-CoV-2 other than S2M11-30 simply cannot induce specific antibody responses (Figure 3 A, B). The KLH-specific serum IgG responses were strongly detected in each peptide-KLH immunization group (Figure 3 C), indicating that immunization operations were in good condition. Next, we analyzed the cross-reactivities of the hyperimmune serum IgG from S2M11-30-KLH, S2M2-30-KLH, and MSM2-19-KLH immunization groups. S2M2-30-KLH-specific serum can strongly and broadly react to peptides S2M11-30, S2M2-20, S1M2-19, as well as S2M2-30 from Omicron BA.1 variant strain (Figure 3 D). Whereas S2M11-30-KLH-specific serum can barely bind any SARS-CoV-2 M peptides other than S2M11-30 itself (Figure 3 E), for which we speculated that its specific serum could ineffectively bind viral M protein ectodomain. MSM2-19-specific serum as a negative control only exhibited binding ability to MSM2-19 (Figure 3 F). In these effective immunization groups, IgG1 was the dominant subclass for the IgG response to M peptides (Figure S3). The immunization groups of S2M2-30-KLH and MSM2-19-KLH demonstrated strong M peptide-specific cellular responses, as shown by the significantly increased IFN-γ-secreting and IL-4-secreting splenocyte populations after re-stimulation with the peptide of S2M2-20, S2M2-30, or MSM2-19 (Figure S4).

**Figure 3.**
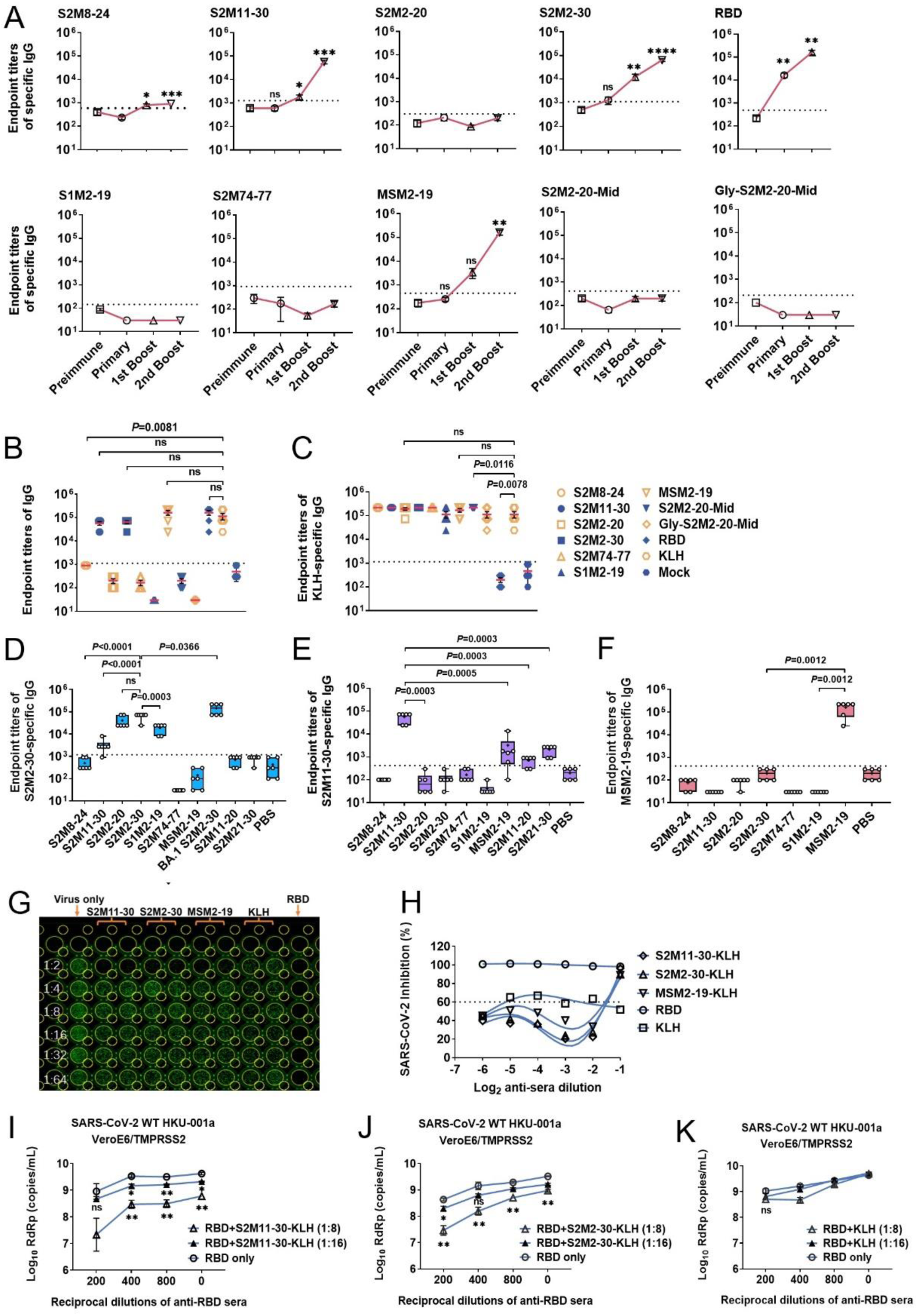
Serological analysis of the immunized BALB/c mice. (A) Titration of serum IgG responses specific to synthetic peptides. The serum samples were collected one day before prime immunization (Preimmune as the negative control) and two weeks post each immunization (Primary, 1st Boost, and 2nd Boost). (B) Comparisons of peptides-specific IgG titers of serum from peptide-KLH-immunized mice, RBD-specific IgG titers of serum from RBD-hFc-immunized mice, KLH-specific IgG titers of serum from unconjugated KLH-immunized mice, and serum from mock-immunized mice. (C) Comparisons of KLH-specific IgG titers of serum from all immunization groups. The mice from the mock-immunization group were intraperitoneally injected with PBS for three times at the same time points as other immunization regimens. (D - F) The analysis of serum cross-reactivities of immune serum collected after the last immunization from groups of (D) S2M2-30-KLH, (E) S2M11-30-KLH, and (F) MSM2-19-KLH. (G, H) Evaluations of serum neutralization activities. (I - K) Microneutralization assays. Analysis of RNA copy numbers of viral RdRp from the wild-type HKU-001a-infected VeroE6-TMPRSS2 cell cultures that were treated with (I) RBD-specific and/or S2M11-30-KLH-specific immune serum, (J) RBD-specific and/or S2M2-30-KLH-specific immune serum, and (K) RBD-specific and/or KLH-specific immune serum, respectively. Data represent Mean ± standard error of the mean (SEM). The dashed lines in the figures represent the Mean + 2 × SEM of the values from negative controls, and values above the dashed lines are regarded as positive results. Statistical significance between compared groups was determined using (A - F) unpaired t-test and (I - K) multiple t-test. P values less than 0.05 are considered statistically significant and ‘ns’ means not significant.

Microneutralization results showed that the hyperimmune serum collected from both S2M2-30-KLH and S2M11-30-KLH immunization groups can neutralize the SARS-CoV-2 wild-type HKU-001a virus *in vitro*. In comparison, WH-1 RBD protein-elicited mice serum exhibited much stronger neutralization activity (Figure 3 G, H). M protein-specific and RBD-specific antibodies often coexist in infected or inactivated vaccine-immunized individuals. We further evaluated the antiviral potencies of the mixed serum with both specificities to M peptide and RBD protein and observed the dosage-dependent synergistic antiviral effects (Figure 3 I, J). In the treatment control groups where RBD-specific serum was absent, the S2M2-30-specific or S2M11-30-specific serum alone could also significantly inhibit the viral replication of SARS-CoV-2 wild-type HKU-001a (Figure 3 I, J), and under no circumstances was ADE detected. No synergistic antiviral effects were detected in the treatment control group where KLH-specific serum was used instead, (Figure 3 K).

Next, we designed an immunization regimen for inducing both RBD-specific and S2M2-30-specific antibody responses in K18-hACE2-transgenic mice, in which mice were immunized with two doses of WH-1 RBD-hFc then followed by three doses of S2M2-30-KLH (Figure S2 B; Table S1). Taking the possible interreaction of immunogens into consideration, RBD-hFc and S2M2-30-KLH were given in separate doses. The IgG titers against RBD or S2M2-30 in the serum samples collected on the 10^th^ day post the 5^th^ dose were determined by using ELISA. Both antigens RBD-hFc and S2M2-30-KLH successfully induced RBD-specific and S2M2-30-specific IgG responses, respectively (Figure 4 A, B). The RBD/S2M2-30-KLH group induced relatively lower RBD-specific IgG titers than RBD group (Figure 4 A), but stronger S2M2-30-specific IgG response than S2M2-30-KLH group (Figure 4 B). The pre-existing RBD-specific immunities in mice seemingly provide a beneficial immune condition conducive to the induction of S2M2-30-specific antibody response while at the expense of being impaired. Anyway, only in the RBD/S2M2-30-KLH immunization group, can the K18-hACE2-transgenic mice acquire robust IgG responses to both RBD and S2M2-30.

**Figure 4.**
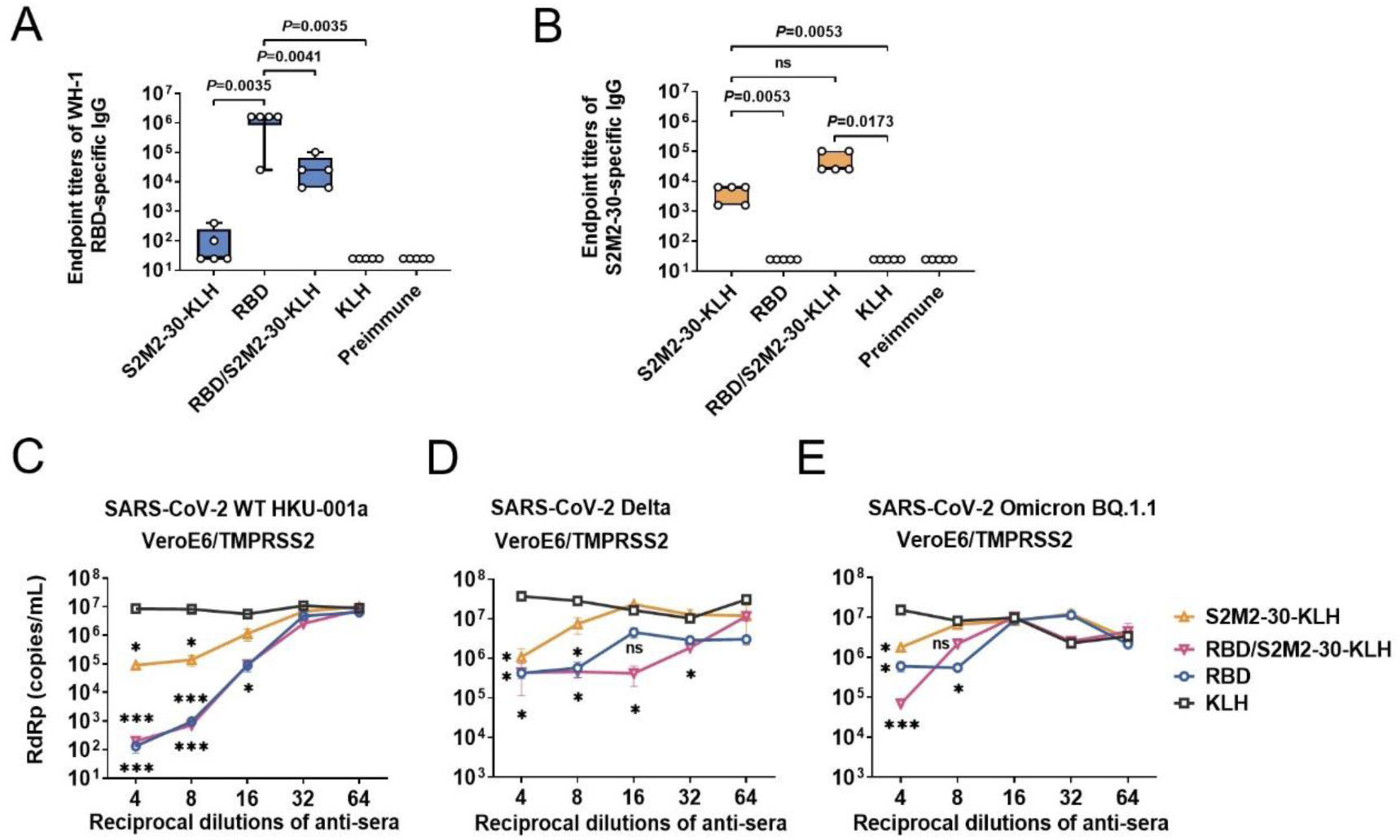
Serological analysis of the immunized K18-hACE2-transgenic mice. (A, B) Titration of serum IgG responses specific to (A) WH-1 RBD protein and (B) S2M2-30 peptide. (C - E) Microneutralization assays. Analysis of RNA copy numbers of viral RdRp from the VeroE6-TMPRSS2 cell cultures infected with (C) wild-type HKU-001a, (D) B.1.617.2/Delta, or (E) Omicron BQ.1.1 viral strain. Data represent Mean ± SEM. Statistical analysis was performed between the values of the KLH group and other groups, and the statistical significance was determined with multiple t-test. The *, **, and *** represent P values less than 0.05, 0.01, and 0.001, respectively. The ‘ns’ indicates not significant.

We found that the RNA copy numbers of SARS-CoV-2 RdRp detected in the cell cultures treated with S2M2-30-specific hyperimmune serum were consistently lower than the KLH-specific serum treatment group (Figure 4 C - E). Be it infections with wild-type HKU-001a, B.1.617.2/Delta, and Omicron BQ.1.1, RBD/S2M2-30-specific hyperimmune serum always exhibited non-inferiority in terms of viral inhibitions to any other specific hyperimmune serum sample, moreover, with certain appropriate dilution factors, was even able to exert stronger antiviral effects (Figure 4 C - E). These results again indicate the advantages of combined antiviral functions over that of serum only specific to either antigen.

### Synergistic immunities conferred heterologous *in vivo* protection

To evaluate immunization potencies in cross-protection, two weeks upon finalizing the immunization regimens, as shown in Figure S2 B, K18-hACE2-transgenic mice were sub-lethally infected with B.1.1.7/Alpha (UK), B.1.1.529/Omicron BA.1, and Omicron BQ.1.1 viruses, respectively. S2M2-30-specific immunity showed *in vivo* cross-protective against the infection with B.1.1.7/Alpha (UK). Two days post intranasal infections of mice with 10^4^ plaque forming unit (PFU) B.1.1.7/Alpha (UK), the RNA copy numbers of viral RdRp detected in the mice lung samples from the groups of S2M2-30-KLH, RBD/S2M2-30-KLH, and RBD were significantly lower than the KLH group, the extents of viral RNA reduction were similar among these protected groups (Figure 5 A). Four days post intranasal infections of mice with 10^5^ PFU B.1.1.529/Omicron BA.1, only in the groups of RBD/S2M2-30-KLH and RBD, were the viral loads in nasal irrigation samples significantly reduced (Figure 5 B). We also recorded and compared the body weight changes and survival rates of immunized mice after sublethal Omicron BA.1 infection (Figure 5 C, D). Upon infection, mice from each group experienced weight loss (Figure 5 C). All mice in RBD/S2M2-30-KLH group survived infection, while the survival rates in the groups of S2M2-30-KLH, RBD, and KLH are 60%, 75%, and 60%, respectively (Figure 5 D). The less protection in the RBD immunization group is due to the large antigenic drift between WH-1 RBD immunogen and viral RBD from Omicron BA.1. Four days post intranasal infections of mice with 10^5^ PFU Omicron BQ.1.1, still only in the groups of RBD/S2M2-30-KLH and RBD, were viral loads in nasal irrigation samples significantly reduced (Figure 5 E). No significant differences in the levels of lung viral RNA were observed among all groups (Figure 5 F).

**Figure 5.**
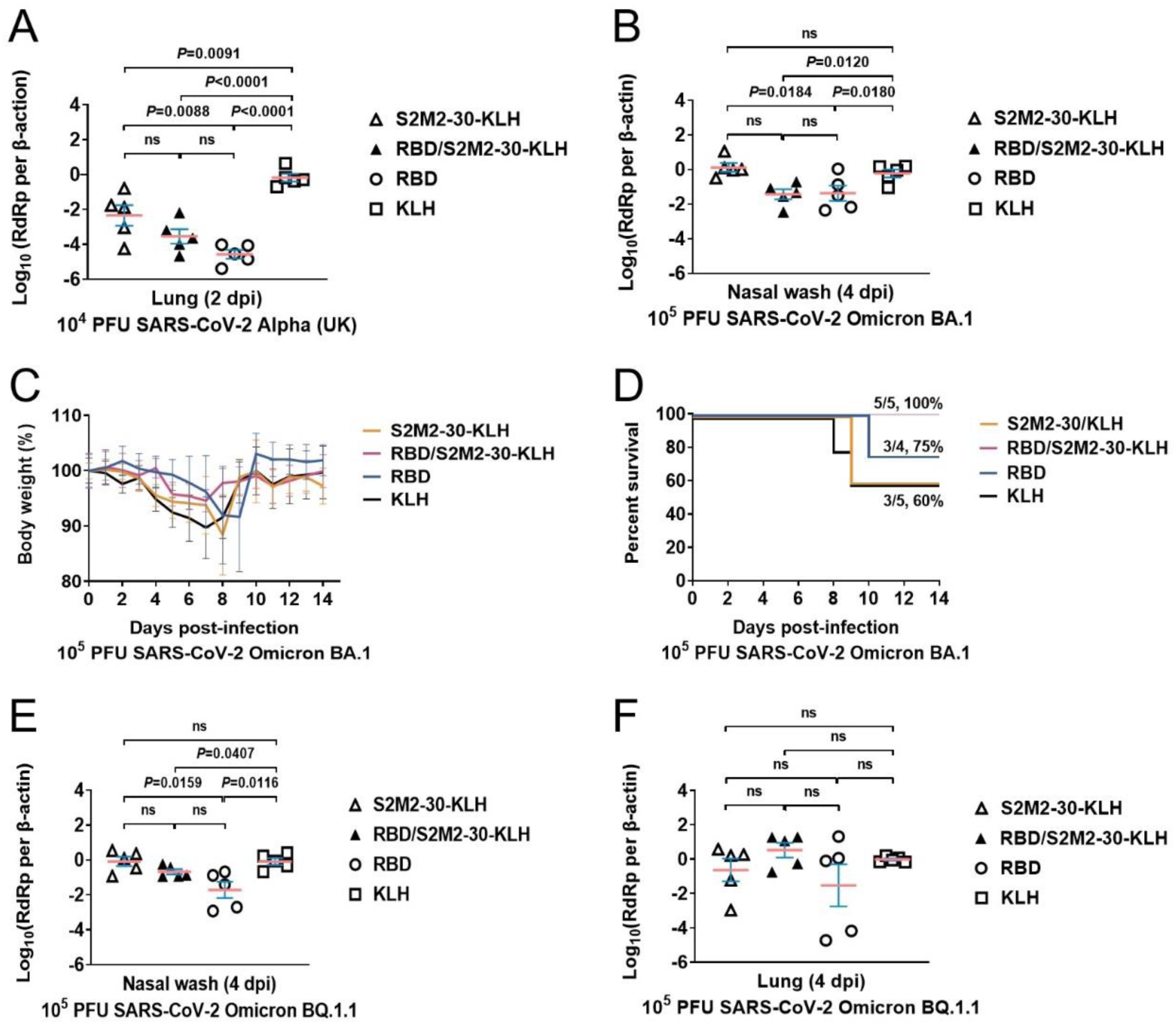
Evaluations of *in vivo* protection against SARS-CoV-2 viruses. (A) Analysis of B.1.1.7/Alpha (UK) viral RdRp gene expression copy number from mice lung homogenate samples by reverse transcription quantitative PCR. The lung tissues from each group were isolated on day two post intranasal infection. (B) Analysis of B.1.1.529/Omicron BA.1 viral RdRp gene expression copy number from mice nasal irrigation samples by reverse transcription quantitative PCR. The nasal irrigation samples from each group were collected on day four post intranasal infection. (C, D) Body weight changes and survival rates of immunization mice were recorded for 14 days after intranasal infection with 10^5^ PFU B.1.1.529/Omicron BA.1. (E, F) Analysis of Omicron BQ.1.1 viral RdRp gene expression copy number from mice nasal irrigation samples and lung homogenate samples by reverse transcription quantitative PCR. The nasal irrigation samples and lung tissues from each group were collected on day four post intranasal infection. All the viral RdRp gene expression copy numbers were normalized to β-actin housekeeping gene expression. Data represent Mean ± SEM. Statistical significance was determined with an unpaired t-test. P values less than 0.05 were regarded as statistically significant. The ‘ns’ indicates not significant.

The protective roles of M peptide-specific antibody responses have been described above. Besides neutralization activities, we also intended to further explore other mechanisms of *in vivo* protectiveness. The successful binding of S2M2-30-specific antibodies to the plasma membrane surface M proteins, which would be regarded as a crucial step in the antibody-dependent cellular antiviral activity, was clearly observed and characterized by using confocal microscopy imaging (Figure 6). We used anti-S2M2-30 serum for immuno-staining SARS-CoV-2-infected A549-TMPRSS2-ACE2 cell culture and confirmed the distribution of the expressed viral M protein on the plasma membrane (Figure 6 A). Besides, the recombinant M protein was also observed on the plasma membrane of HEK293T cell culture that was transfected with a eucaryotic plasmid encoding the recombinant fusion protein comprising M protein and green fluorescent protein (Figure 6 B). At last, the immuno-fluorescent staining showed the specific bindings of anti-S2M2-30 serum antibodies to HEK293T cells expressing recombinant M protein (Figure 6 C).

**Figure 6.**
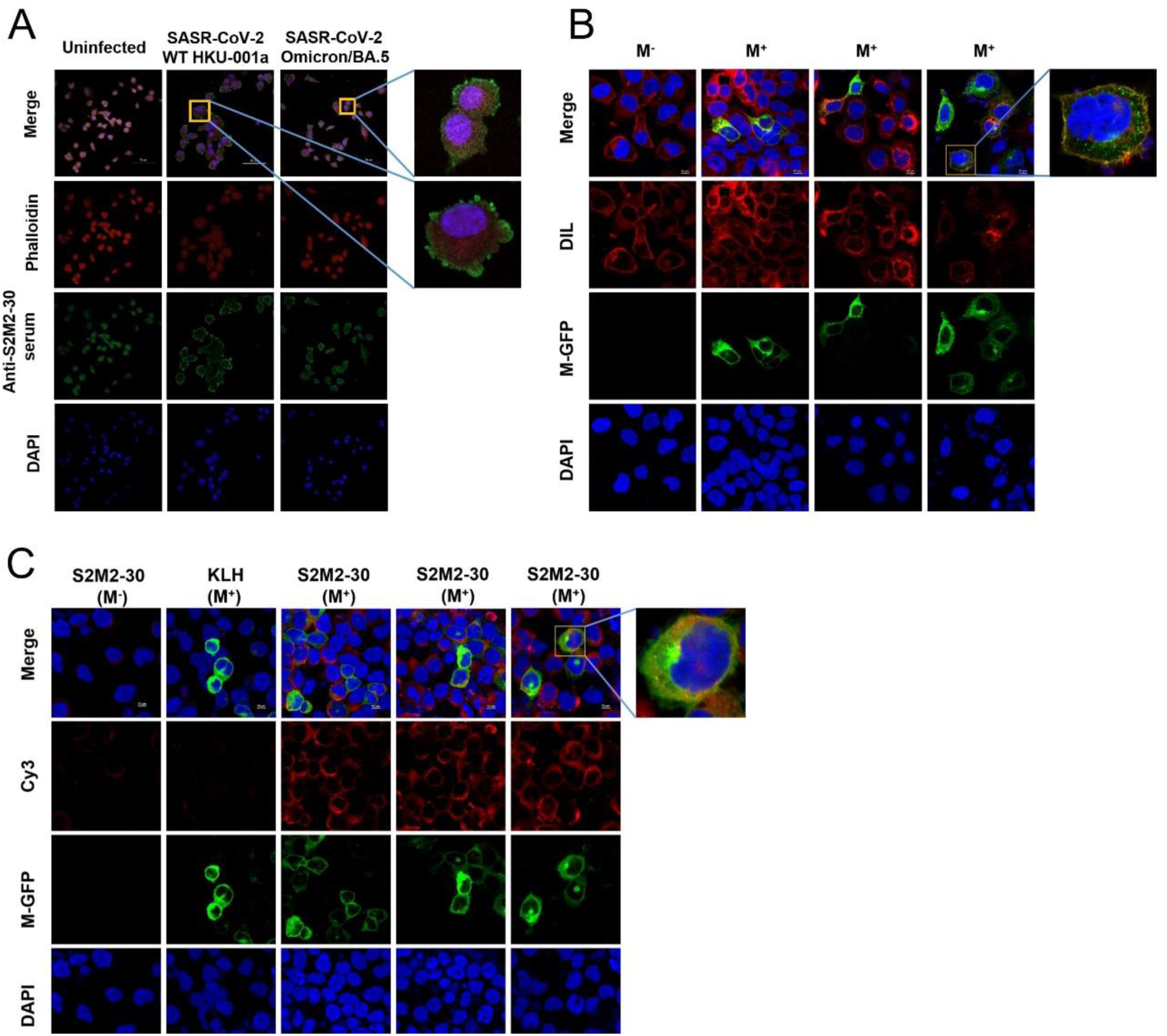
Immunofluorescence analysis. (A) Immunofluorescent staining analysis of the SARS-CoV-2 virus-infected VeroE6-TMPRSS2 cells with S2M2-30-specific hyperimmune serum. The cells were infected with wild-type HKU-001a and Omicron/BA.5, respectively, and the uninfected cells were used as negative controls for immunostaining. The phalloidin-iFluor dye and DAPI were used for staining the F-actin and nucleus in cells, respectively. (B) Confocal fluorescence microscopy analysis of the recombinant GFP-M fusion protein expression in the transfected HEK293T cells. The DIL dye and DAPI were used for staining the cell membrane and cell nucleus, respectively. (C) Immunofluorescent staining analysis of the transfected HEK293T cells over-expressing GFP-M fusion protein with S2M2-30-specific hyperimmune serum. Fluorescently labeled secondary antibody Cy3 Goat Anti-Mouse IgG H&L was used for the signal amplification of primary mouse antibody detection.

## Discussion

To our initial understanding, the antibodies specific to the M protein ectodomain rather than transmembrane and cytosolic domain potentially impede viral infection. This study achieves the goal to solely evaluate the functions of the immunities specific to the N-terminal peptides of M protein, and reveals their cross-protective roles in combating authentic SARS-CoV-2 viruses *in vitro* and *in vivo*. So far as we know, the protection mechanisms of S2M2-30-specific immunity involve viral neutralization and also antibody-dependent cell-mediated cytotoxicity as well. We found no experimental evidence of the ADE effect of its specific immunities to the N-terminal peptides. High seroconversion rate to the N-terminal peptides were detected in the serum samples collected from convalescent patients and vaccinees receiving three shots with inactivated SARS-CoV-2 vaccines (Figure 2). We strongly believe that the detected specific human immunities to M protein ectodomain are of benefit to the *in vivo* resistance to SARS-CoV-2 infections.

Our immunological investigations offer a few tips to facilitate a better understanding of the protective peptide immunogen designs. Circular dichroism analysis suggested a relatively more random coil structure in S2M2-20 rather than in S2M11-30 and S2M2-30 (Figure 1 E). The flexible loop in the S2M2-20 peptide is inconducive to the induction of specific antibody response, which could be a reason behind the less efficient seroconversion to the N-terminal peptide in many individuals ever receiving inactivated vaccines. In the development of M peptide-based vaccines, the shorter peptides that are closer to the extreme N-terminal of M protein, such as S2M2-20, S2M8-24, and S1M2-19, were found in a lack of mouse B-cell epitopes and difficult in efficiently hooking B-cell receptor (BCR) and eliciting antibody responses (Figure 3 A). In contrast, the S2M2-30 peptide in which at least one complete B-cell epitope exists effectively elicited protective antibody responses when conjugated with KLH carrier. Reproducing the native conformation of viral peptides in KLH conjugates is extremely hard. The good protectiveness of S2M2-30-specific antibodies indirectly illustrates a fact that the N-terminal peptide in part presents flexible conformation, which is consistent with the results of our circular dichroism analysis. Therefore, without much rigid conformation constraints, vaccinations with the S2M2-30 presented by many immunogenic carriers could readily induce protective antibody responses.

Even though the S2M21-30 peptide of M protein is generally regarded in the transmembrane spanning region (Figure 1 A), interestingly, this fragment of amino acids is quite necessary for forming the complete B-cell epitope of S2M2-30 peptide and efficiently inducing M protein ectodomain-specific antibodies (Figure 3 A, B). The cryo-electron microscopy analysis illustrated the compact architecture of the tandemly arranged viral M protein oligomers ^9^, implying no room in between protomers left for the lipid layer. We infer that S2M21-30 fragments on the SARS-CoV-2 virion surface or cellular plasma membrane are most likely not buried in the lipid layer and can be readily recognized by BCR. For now, we cannot rule out the possibility that M protein ectodomain induces specific antibody responses in a quaternary structure-dependent manner.

The neutralization activities of S2M2-30-specific serum was obviously detected in the microneutralization assays (Figure 3 G, H, J; Figure 4). Tomogram of SARS-CoV-2 virions illustrated the distribution of S proteins nearest-neighbor distances on virions’ surface with an average of 23.6 nm ^13^. The considerable space most probably allows massive antibodies reach and bind to the M protein ectodomains beneath S proteins on the virion surface. The M protein ectodomain-specific antibodies bound on the virion surface could not only mitigate the viral attachment to host cells but also interfere the post-fusion structure change of S protein. We speculate that the mutations D3G/N and Q19E in the SARS-CoV-2 Omicron M protein (Figure 1 C) might slightly impair the specific binding of S2M2-30-specific antibodies, which weakens *in vitro* serum neutralization activities against Omicron BQ.1.1 (Figure 4 C - E) and *in vivo* cross-protection against Omicron BA.1 and BQ.1.1 virus infections (Figure 5).

Serum IgG responses to SARS-CoV-2 M protein ectodomain were general among human populations who were exposed to M protein antigen and relatively stronger in convalescent patients (Figure 2 A). We found that once combined with RBD-specific serum antibodies it more likely than not exerts synergistic antiviral effect with broader and stronger cross-neutralization activities (Figure 3 I - K, and Figure 4 C -E). Our experiment results hint stronger *in vivo* cross-protection in humans could be provided by immunization with a blend of conserved antigens, thus the M protein ectodomain-based immunogen holds promise as a supplemental for such a universal sarbecovirus vaccine. In the next step, it is necessary to perform in-depth profiling of specific antibody repertoire and clarify the distribution of human B-cell epitopes in the M protein ectodomain for the sake of the universal immunogen design.

## Methods and Materials

### Human subjects

In this study, we recruited vaccinees from a prospective cohort at the Third Affiliated Hospital of Sun Yat-sen University in Guangzhou, China, all participants received the inactivated SARS-CoV-2 vaccine BBIBP-CorV (BBIBP-CorV, Sinopharm, Beijing) or CoronaVac (Sinovac Life Sciences, Beijing, China) on day 0, 28 and 270. Serum samples of vaccinees who received two doses of inactivated vaccines (n=58), and serum samples of vaccinees who received three doses of inactivated vaccines (n=30) were collected 14 days after the last immunization. Convalescent serum samples of COVID-19 patients (n=15) were collected 14 days after release from hospitalization. Our study also included serum samples of child outpatients (n=28) in the middle of the year 2019 for negative control use in serological tests. All serum samples were heat-inactivated at 56°C for 30 min before use.

### Cell lines and SARS-CoV-2 strains

VeroE6 cell line with TMPRSS2 overexpression was purchased from the Japanese Collection of Research Bioresources (JCRB) Cell Bank and maintained in Dulbecco’s modified Eagle’s medium (DMEM) (Gibco, Cat.#11965092) supplemented with 10% heat-inactivated fetal bovine serum (FBS) (Gibco, Cat.#16140071), 100 U/ml penicillin and 100 μg/ml streptomycin (P/S) (Gibco, Cat.#15140122) and 1mg/ml G-418 (MCE, Cat.#HY-17561). A549 with TMPRSS2 and human ACE2 overexpression cell line that was used in immunostaining was purchased from InvivoGen company (Cat.#a549-hace2tpsa) and maintained in DMEM supplemented with 0.5 μg/ml of Puromycin (Thermofisher, Cat.#A1113803) and 300 μg/ml of Hygromycin (Thermofisher, Cat.#10687010), 10% FBS, P/S. The following authentic SARS-CoV-2 viruses including wild-type HKU-001.a (GenBank: MT230904), B.1.1.7/Alpha (UK) (GenBank: OM212469), B.1.617.2/Delta (GenBank: OM212471), B.1.1.529/Omicron BA.1 (GenBank: OM212472), Omicron BA.5 (GISAID: EPI_ISL_13777658), Omicron BQ.1.1 (GISAID: EPI_ISL_16342297) viruses were isolated from nasopharyngeal swabs of laboratory confirmed COVID-19 patients and were used in this study. All the viruses were sub-cultured and tittered by plaque assay using VeroE6-TMPRSS2 cells. Each virus sample was aliquoted and stored at -80 °C for further use. All authentic SARS-CoV-2 related experiments were performed in Biosafety Level 3 (BSL-3) Facility at the Department of Microbiology, The University of Hong Kong under the Standard Operation Procedures (SOPs).

### Laboratory mouse models

Six-to-eight-week-old female BALB/c mice were obtained from the Hunan SJA Laboratory Animal Co., Ltd. and maintained with access to food and water ad libitum in a clean room kept at 22 ± 2 °C and 50 ± 5% humidity, under a 12 h-12 h light-dark cycle. All animal experiments were performed in accordance with the China Public Health Service Guide for the Care and Use of Laboratory Animals. Experiments involving mice and protocols were approved by the Institutional Animal Care and Use Committee from College of Biology, Hunan University (Approval Code: HNUBIO202202002).

Six-to-eight-week-old K18-hACE2 transgenic mice were purchased from The Jackson Laboratory (US) and kept in cages with individual ventilation under 65% humidity and an ambient temperature of 21–23 °C and a 12 h–12 h day–night cycle for housing and husbandry in Centre for Comparative Medicine Research (CCMR) of The University of Hong Kong. All mice were housed and bred in our BSL-3 facility ^18^ and given access to standard pellet feed and water ad libitum.

### M protein structure prediction with AlphaFold-2

Sequences for coronavirus M proteins were obtained from UniProt: P59596, VME1_SARS; P0DTC5, VME1_SARS2 and K9N7A1, VME1_MERS1. The amino acid sequences of coronavirus M proteins were submitted to Google Colab (https://colab.research.google.com/github/sokrypton/ColabFold/blob/main/beta/AlphaFold2_advanced.ipynb) and the structures of full-length coronavirus M proteins were predicted by running AlphaFold2_advanced.ipynb. The structure prediction process was described in the AlphaFold2_advanced (Google Colab), consisting of five steps: MSA construction, template search, inference with five models, model ranking based on mean pLDDT, and constrained relaxation of the predicted structures. The highest-ranked models were selected for visualization with ChimeraX software (UCSF, https://www.cgl.ucsf.edu/chimerax/).

### Peptides synthesis

The set of peptides listed in Table 1 was synthesized by a standard solid-phase Fmoc (9-fluorenyl methoxy carbonyl) method in Shanghai Top-Peptide Biochem Co., Ltd. Peptide was purified to homogeneity (purity of >95%) by high-performance liquid chromatography and identified by laser desorption mass spectrometry. A monosaccharide N-acetylglucosamine was linked to the fifth asparagine of Gly-S2M2-20-Mid peptide.

### Circular dichroism (CD) spectroscopy

The secondary structures of the synthetic peptides in solid state were confirmed by far UV CD spectroscopy using a JASCO J-815 spectrometer (JASCO Analytical Instruments, Easton, MD). The lyophilized peptide powders were spread on 2-cm diameter cylindrical quartz glass, and the spectra of samples were recorded over a wavelength range of 190-260 nm at a scan speed of 10 nm/min and a time constant of 0.5 s. An average of three scans was reported. The data of CD spectra were analyzed by CDNN CD spectral deconvolution software package (version 2.1, Applied Photophysics Ltd., Leatherhead, UK), and the proportion of each secondary structure (α-helix, β-antiparallel, β-turn, unordered random coli) was calculated by running “deconvolute”. The final spectra were expressed as molar circular dichroism Δε (L·mol^−1^·cm^−1^) per residue or molar absorption coefficient ε (L·mol^−1^·cm^−1^) per residue.

### Preparation of peptide-KLH conjugates

The conjugations of peptides to the KLH protein carrier were performed according to the manufacturer instructions of ReadiLink™ KLH Conjugation Kit (AAT Bioquest, Cat.#5502). In brief, 1 mg soluble peptide was dissolved in 0.5 ml PBS and mixed with 1 mg of KLH in 0.5 ml PBS. Those peptides poorly solubilized in PBS, including S2M11-30, S2M2-30, and MSM2-19, were dissolved in 0.5 ml PBS containing 5% dimethyl sulfoxide (DMSO, Solarbio, Cat.#D8371) and mixed with KLH. 50 μl 1% (w:v) of 1-ethyl 1-3- [dimethylaminopropyl] carbodiimide hydrochloride (EDC) was immediately added to the mixture of peptide and KLH, then gently mixed and incubated at room temperature for 2 h. Spin desalting columns (7K MWCO) were used to purify the conjugates and to remove the non-reacted crosslinker. The collected conjugate samples were sterilely filtered by using a 0.22 μm filter membrane, then stored at -80°C.

### Mice immunization and infection experiments

BALB/c mice were used to assess immunogenicity of peptide-KLH conjugates. The immunization regimen is depicted in Figure S2 A. Each group (n = 6) was immunized intraperitoneally with peptides-KLH (10 μg/mouse) emulsified in Incomplete Freund’s Adjuvant (Sigma, Cat.#F5506), and a control group (n = 6) was immunized with unconjugated KLH. The first and second booster immunizations were performed with the same dose of antigens emulsified in Incomplete Freund’s Adjuvant on day 21 and day 42 after initial immunization, respectively. The SARS-CoV-2 WH-1 RBD-hFc recombinant protein (Sino Biological Inc., Cat.#40592-V02H) immunization group was immunized twice with RBD-hFc, 10 μg per immunization dose. The mice were bled one day before the prime immunization and two weeks after each immunization for antibody titration using ELISA. All groups of mice were sacrificed by cervical dislocation on day 63, then blood samples and spleens were collected for subsequent experiments.

Six-eight-week-old male K18-hACE2 transgenic mice were used for the SARS-CoV-2 challenge experiments ^18^ and the immunization regimens were depicted in Figure S2 B. The mice were firstly immunized with 2 doses of RBD-hFc or just sterile PBS for mock immunization via intramuscular injection at 14 days intervals. After two immunizations with RBD-hFc or PBS, the mice were immunized with additional 3 doses of S2M2-30-KLH conjugate or unconjugated KLH at 14 days intervals. Blood was taken one day before prime immunization and 10 days post the second, fourth, and fifth immunizations. Two weeks after the last immunization, the mice were anesthetized with Ketamine (200 mg/kg) (purchased from CCMR, HKU) and Xylazine (10 mg/kg) (purchased from CCMR, HKU), followed by intranasal inoculation of SARS-CoV-2 viruses (10^4^ PFU inocula for B.1.1.7/Alpha (UK) viral challenge; 10^5^ PFU inocula for both Omicron BA.1 and BQ.1.1 viral challenges). Body weight and survival rate were recorded daily for 14 days after Omicron BA.1 infection. Nasal irrigation and lung samples were collected for viral RNA copy detection. All SARS-CoV-2 infection experiments were performed in the BSL-3 facility and were approved by the Committee on the Use of Live Animals in Teaching and Research (CULATR) of The University of Hong Kong.

The infected cell culture or animal samples were lysed by using RLT buffer (Qiagen, Cat.#79216), followed by viral RNA extraction using RNeasy Kit (Qiagen, Cat.# 74004) according to the manufacturer’s protocol. The viral copies were quantitated by reverse transcription quantitative PCR following the handbook (Takara, Cat.#RR086A). The copy numbers of reverse-transcripts from cell cultures and mice organ samples were normalized with the expression level housekeeping gene of beta-actin by using the 2(-Delta Delta C(T)) method^19^. The primers used in this study targeting SARS-CoV-2 RNA-dependent RNA Polymerase (RdRp) gene include Forward primer: 5’- CGCATACAGTCTTCAGGCT-3’ and Reverse primer: 5’- GTGTGATGTTGAWATGACATGGTC-3’, and the primers targeting beta-actin gene include Forward primer: 5’-ACGGCCAGGTCATCACTATTG-3’ and Reverse primer: 5’-CAAGAAGGAAGGCTGGAAAAG-3’.

### Serological analysis

BALB/c mice were submandibular bled 1 day before prime immunization, and 14 days after each immunization. K18-hACE2-trangenic mice were bled 1 day before prime immunization, and 10 days after the 2^nd^, 4^th^, and 5^th^ immunizations. Titers of peptide-specific and RBD-specific serum IgG were determined by enzyme-linked immunosorbent assay (ELISA) using 96-well immuno-plates (JET BIOFIL, Cat.#FEP100096) coated overnight with 100 ng per well of peptides (Table 1) or purified WH-1 RBD protein (WH-1 RBD (Arg319 - Phe541), GenBank: QHD43416.1). Prior to incubation with serum samples, the immuno-plates were blocked with 3% bovine serum albumin (BSA, Sigma, Cat.#0336-50ML) in 1 × PBS supplemented with 0.1% (v:v) Tween-20 (PBST) at 37 °C for 1 h. After blocking, the immuno-plates were incubated with three-fold serially diluted serum samples starting from 1:100 to 1:218700 at 37 °C for 1 h. Immuno-plates were then washed three times with 0.1% PBST and incubated with 0.2 μg/ml goat anti-mouse IgG conjugated with horseradish peroxidase (HRP) (ABclonal Technology, Cat.#AS003) in blocking buffer at 37 °C for 1 h. Immuno-plates were washed four times and developed by using TMB substrate (Solarbio, Cat.#K8160) at room temperature for 10 min. The reaction was stopped with 1 M sulfuric acid and the absorbance was measured at 450 nm.

The cut-off for seropositivity was set as the mean + 2 SD value of the preimmune serum control group and the level was indicated as the dashed line. The HRP-conjugated goat anti-mouse IgG1 (ABclonal Technology, Cat.#AS066), the HRP-conjugated goat anti-mouse IgG2a (ABclonal Technology, Cat.#AS065), or the HRP-conjugated goat anti-mouse IgM (Sangon Biotech, Cat.#D110103-0100) was used in ELISA, for titrating specific serum IgG1, IgG2a, IgM antibodies.

The ELISA experiment procedures for titrating specific antibodies of human serum samples were slightly modified. In brief, High-binding 96 well ELISA plates (Thermo, Cat.#442404) were coated with antigen 50 μl per well, 2 μg/ml, in 1 × carbonate-bicarbonate buffer at 37 °C overnight until the coating were dry. In the following day, the wells were blocked with 300 µl blocking buffer, containing 5% skim milk powder (BBI, Cat.#A600669-0250) in PBST, at room temperature for 1 h. Human blood serum was diluted 300 and 900 times for binding to antigen and incubated at room temperature for 2 h. After washing 3 times with washing buffer, the secondary anti-human IgG antibody conjugated to HRP (Jackson ImmmunoResearch, Cat.#109-035-088) diluted 1:2000 in PBS, and incubated at room temperature for 1 h. Immuno-plates were washed four times and developed by using TMB substrate at room temperature for 10 min. The reaction was stopped with 1 M sulfuric acid, absorbance was measured at 450 nm on a microplate spectrophotometer (BioTek) and the OD values were used to calculate AUC using GraphPad Prism 8.0.

### Microneutralization assays

In the microneutralization assays, serial-fold diluted immune serum samples were pre-incubated with SARS-CoV-2 viruses at 37 °C for 1 h. The serum-virus mixtures were then inoculated in the confluent VeroE6-TMPRSS2 cell culture for 1 h. The cell cultures were washed twice by using sterile PBS. Then, the cell cultures were maintained with serum diluents at 37 °C for 48 h. The culture supernatants were harvested for reverse transcription quantitative PCR detection of viral load. For the evaluations of synergistic antiviral effects of the mixed serum, serially diluted (1:200, 400, or 800) RBD-specific immune serum samples were mixed with the serially diluted (1:8 or 16) peptide-KLH-specific or KLH-specific immune serum samples. For evaluations of the antiviral effects of serum from immunized K18-hACE2-transgenic mice, the serum samples were 3-fold serially diluted for specific antibody titrations.

We also used the method of immune-fluorescent staining HKU-001a-infected Vero-E6-TMPRSS2 cells for the evaluation of serum neutralization activities. cells were seeded into 96-well cell plates (Corning, Cat.#3904) at a density of 20,000 cells/well and incubated at 37 °C under 5% CO_2_ overnight. The serum samples were 2-fold serially diluted with an initial dilution factor of 1:2, then each serum diluent sample was mixed 10^3^ PFU/ml SARS-CoV-2 HKU-001a virus with an equal volume and incubated at 37 °C for 1 h. Virus-serum mixtures were added to cell culture plates for infections, the virus samples without mixing with serum were used as positive controls for infection. After 24 h incubation, the supernatant was removed and the cells were fixed with 4% paraformaldehyde at room temperature for 15 min. The cell cultures were washed three times with PBS and permeabilized with 100 μl 0.5% TritonX-100 (Sigma, Cat.#93443) at room temperature for 15 min. After washing three times with PBS, the cells were incubated with 1:4000 fold diluted rabbit anti-SARS-CoV-2 N protein-specific IgG (Abcam, Cat.#ab271180) at room temperature for 1 h. After washing three times with PBS, 1:100 fold diluted Goat anti-Rabbit IgG (H+L) Cross-Adsorbed Secondary Antibody (Alexa Fluor™ 488, ThermoFisher, Cat.#R37116) was then added and incubated at room temperature for 1 h. The fluorescence intensity of each well was read using Sapphire Biomolecular Imager (Azure Biosystems). Quantitative analysis of the microneutralization experiment was done with ReadPlate 3.0 in ImageJ software.

### Immunofluorescent staining analysis

HEK293T cell line was used in plasmid transfection experiment for the expression of SARS-CoV-2 full-length M protein. The pre-seeded HEK293T cells at the density of 1×10^5^ cells per well were grown at 37 °C overnight on the glass coverslips in 96-well cell culture plates (JET BIOFIL, Cat#TCP011096). Then, each well was transfected with 2 μg eucaryotic recombinant plasmid pCMV3-C-GFPSpark (C-GFPSpark tag, Sino Biological Inc., Cat#VG40608-ACG) encoding the SARS-CoV-2 full-length membrane glycoprotein fused to a green fluorescent protein (GFP). 6 μl Lipofectamine 3000 (Thermo Fisher, Cat#L3000001) was used for encapsulating 2 μg plasmid, the lipid complex was wise-dripped into each well and incubated at 37 °C, 5% CO_2_ for 6 h. Then refresh the culture medium with DMEM supplemental with 10% FBS and the cells were incubated at 37 °C, 5% CO_2_ for an additional 24 h. The successful expression of the GFP-M fusion protein was confirmed by observing the green fluorescence using the inverted phase contrast fluorescence microscopy (Nikon, Cat.#TS2R-FL, Japan). The 1,1’-dioctadecyl-3,3,3’,3’-tetramethylindocarbocyanine perchlorate (DIL, Solarbio, Cat.#D8700) and 2- (4-Amidinophenyl)-6-indolecarbamidine dihydrochloride (DAPI, Cat.#D8700) were used for staining cell membrane and cell nucleus, respectively.

For immunostaining of the transfected cells, the HEK293T cells over-expressing GFP-M fusion protein were fixed using 4% paraformaldehyde (Solarbio, Cat.#P1110) in tris buffered saline (TBS) at 4°C for 15 min, permeabilized using 0.2% Triton X-100 (Sigma, Cat#93443) in TBS at room temperature for 20 min, blocked using 10% normal goat serum (Solarbio, Cat#SL038) at room temperature for 1 h, and incubated with diluted S2M2-30-KLH antiserum (1:100) at 4 °C overnight. After washing with PBS three times, 5 min each time, the cells were incubated with Cy3 Goat Anti-Mouse IgG H&L (ABclonal, Cat#AS008) at room temperature for 1 h. After washing with PBS for 3 times, the cells were stained with 2 μg/ml DAPI at room temperature for 5 min. The stained cells on the coverslips were imaged using a laser scanning confocal microscope (ECLIPSE, Nikon, Japan).

For immunostaining of the infected cell cultures, pre-seeded A549-TMPRSS2-ACE cells were infected with SARS-CoV-2 wild-type HKU-001a or Omicron/BA.5 variant with MOI of 2. At 2 hour-post-infection, cell culture supernatant was discarded, and cells were fixed using 4% formalin at room temperature for 15 min and blocked with 10% skimmed milk for 1 h. The cells were incubated with S2M2-30-KLH-specific immune serum at room temperature for 1 h, then were washed 3 times using PBST, followed by incubation of secondary antibody Goat Anti-Mouse IgG H&L (Alexa Fluor 488, Abcam, Cat.#ab150113). The phalloidin-iFluor dye (Abcam, Cat.#ab176759) and DAPI were used for staining the F-actin and nucleus in cells, respectively. The images were taken by Carl Zeiss LSM880 system Confocal Laser Scanning Microscope (Dublin, CA, USA).

### The enzyme-linked immunospot (ELISPOT) assay

Mice were sacrificed 3 weeks post the last immunization and soaked in 75% ethanol for a while before spleen isolation. The spleen organs were aseptically excised and isolated and then were cut into small pieces in cell culture dishes using sterilized ophthalmic scissors and ground using the flat end of a syringe piston, then the samples were transferred into 40 μm cell strainer and washed with 5 - 10 ml RPMI 1640 medium (Gibco, Cat.#11875119). The suspension of splenocyte single cells was immediately transferred to a 15 ml sterile centrifugation tube and centrifuged at 300 × g at 4 °C for 10 min. After centrifugation, lymphocytes were isolated by Mouse Spleen Lymphocyte Separation Kit (Solarbio, Cat.#P8860). Lymphocyte pellets were harvested after the centrifugation at 250 × g at 4 °C for 10 min. The cells were resuspended in RPMI 1640 medium at the cell density of around 10^8^ cells per ml for subsequent seeding plates. Interferon (IFN)-γ-positive cells were detected using a mouse IFN-γ ELISPOT Kit (BD, Cat.#551083, U.S.), and interleukin (IL)-4-positive cells were detected using a mouse IL-4 ELISPOT Kit (DAKEWE, Cat.#2210403). Briefly, coated plates were washed three times with PBS and blocked with RPMI-1640 medium containing 10% FBS (Life Technologies, Cat.#A0336-50ML) at 37 °C for one and a half hours. Freshly prepared single-cell suspensions were seeded in the ELISPOT immuno-plates at the cell density of 2 × 10^6^ cells per well and re-stimulated with SARS-CoV-2 or MERS M peptides, including S2M2-20, S2M2-30, or MSM2-19 with 1 µg per well, at 37 °C for 24 h, Concanavalin A (ConA, Sigma, Cat.#C2272) at a final concentration 2 μg/ml was used as a positive control for reactivating T lymphocytes. After restimulation with peptides, cells were lysed with ice-cold deionized water. Then the plates were incubated with anti-mouse IFN-γ or IL-4 detection antibody (at 1:100 dilution) at room temperature for 2 h, afterward with 100 µl each well of streptavidin-HRP (at 1:100 dilution) for 1 h. Spots corresponding to antigen-specific antibody-secreting cells were developed using BD ELISPOT AEC Substrate Set (BD, Cat.#551951) at room temperature for 15 min, and then the immuno-plates were washed twice with deionized water to stop the reaction. Spots were counted using an ELISPOT automatic plate reader (AID Elispot Reader, AID, Germany).

### Statistical Analysis

All data in comparison were analyzed with GraphPad Prism 9 software v9.1.1. Statistical significance between two experimental groups in comparison was analyzed using unpaired two-tailed Student’s t-tests (Figure 2 A, Figure 3 A - F, Figure 4 A - B, Figure 5 A - B, Figure 5 E - F, Figure S1 B, Figure S3, and Figure S4). Comparisons among three or more experimental groups were analyzed using two-way ANOVA with Tukey’s multiple-comparison test (Figure 2 B). Comparisons of SARS-CoV2 RdRp RNA copies of different groups from microneutralization assays were performed using multiple t-tests. (Figure 3 I - K and Figure 4 C - E). The statistical significance of survival rates between groups in viral challenge experiments was analyzed using the log-rank (Mantel– Cox) test (Figure 5 D). Differences were considered to be statistically significant when *P* < 0.05. The ‘ns’ indicate no significant difference between groups in comparison.

## Ethics statement

The proposal for performing the BALB/c mice immunization experiments in an animal laboratory from Hunan University was reviewed and approved by the Animal Ethics Committee from the College of Biology, Hunan University (Ethics Approval Code: HNUBIO202202001).

All SARS-CoV-2 infection experiments involving animals were performed in the BSL-3 facility and were approved by the Committee on the Use of Live Animals in Teaching and Research (CULATR) of The University of Hong Kong (Ethical number: CULATR 5440-20 and 5370-20).

The serological analysis of all human serum samples in this study was approved by the Research Ethics Committee of the School of Public Health (Shenzhen), Sun Yat-sen University, China, and all patients signed Informed Consent Forms. SARS-CoV-2 nasal swabs reverse transcription PCR tests were used to confirm that all COVID-19 convalescent patients were negative for SARS-CoV-2 at the time of blood collection. The blood collection time of all COVID-19 convalescent patients was 14 days after discharge from the hospital.

## Acknowledgments

This study was supported by the National Natural Science Foundation of China (Project Approval Number: 81971566) and Hunan Province Key R&D Plan (Project Approval Number: 2021SK2030) to Lei Deng; Guangdong Natural Science Foundation (2022A1515010099 and 2023A1515012907), National Key Research and Development Program of China (2021YFC0866100), and Theme-Based Research Scheme (T11-709/21-N) of the Research Grants Council and Hong Kong Special Administrative Region to Shuofeng Yuan; the National Natural Science Foundation of China (92169104 and 31970881), Shenzhen Science and Technology Program (JCYJ20200109142438111, KQTD20200820145822023, RCJC20210706092009004, and GXWD20201231165807008) grants to Yaoqing Chen, Double-First Class Construction Funds of Hunan University (521119400156) to Xingyi Ge and Lei Deng.

## Author contributions

Lei Deng conceived the project. Yibo Tang, Lei Deng, Shuofeng Yuan, Yaoqing Chen, and Xingyi Ge participated in the experimental design work. Yibo Tang performed the circular dichroism analysis of synthetic peptides, prepared the peptide-KLH conjugate immunogens, conducted computational modeling of M protein structures, performed the BALB/c mice immunization experiments, the serological analysis and cellular responses thereof, and immunostaining of transfected HEK293T cells. Kaiming Tang, Cuiting Luo, and Hehe Cao performed the K18-hACE2 transgenic mice immunization and infection experiments and cell culture infection experiments in the BSL-3 lab from The University of Hong Kong. Yunqi Hu, Wanyu Luo, and Yibo Tang independently analyzed the human serum antibody titers specific to M ectodomain peptides and RBD protein, at the School of Public Health (Shenzhen), Sun Yat-sen University and the College of Biology from Hunan University, respectively. Yibo Tang, Kaiming Tang, Cuiting Luo, Yunqi Hu, Zi-Wei Ye, Dejian Liu, Cuicui Liu, Rang Wang, Xingyi Ge, Yaoqing Chen, Shuofeng Yuan, and Lei Deng participated in collecting and/or analyzing experimental data. Yibo Tang made the figures and tables. Lei Deng and Yibo Tang drafted the manuscript and all authors proofread the manuscript. Lei Deng, Shuofeng Yuan, and Yaoqing Chen are accountable for the accuracy and integrity of this work.

## Supplementary Information

**Figure S1.**
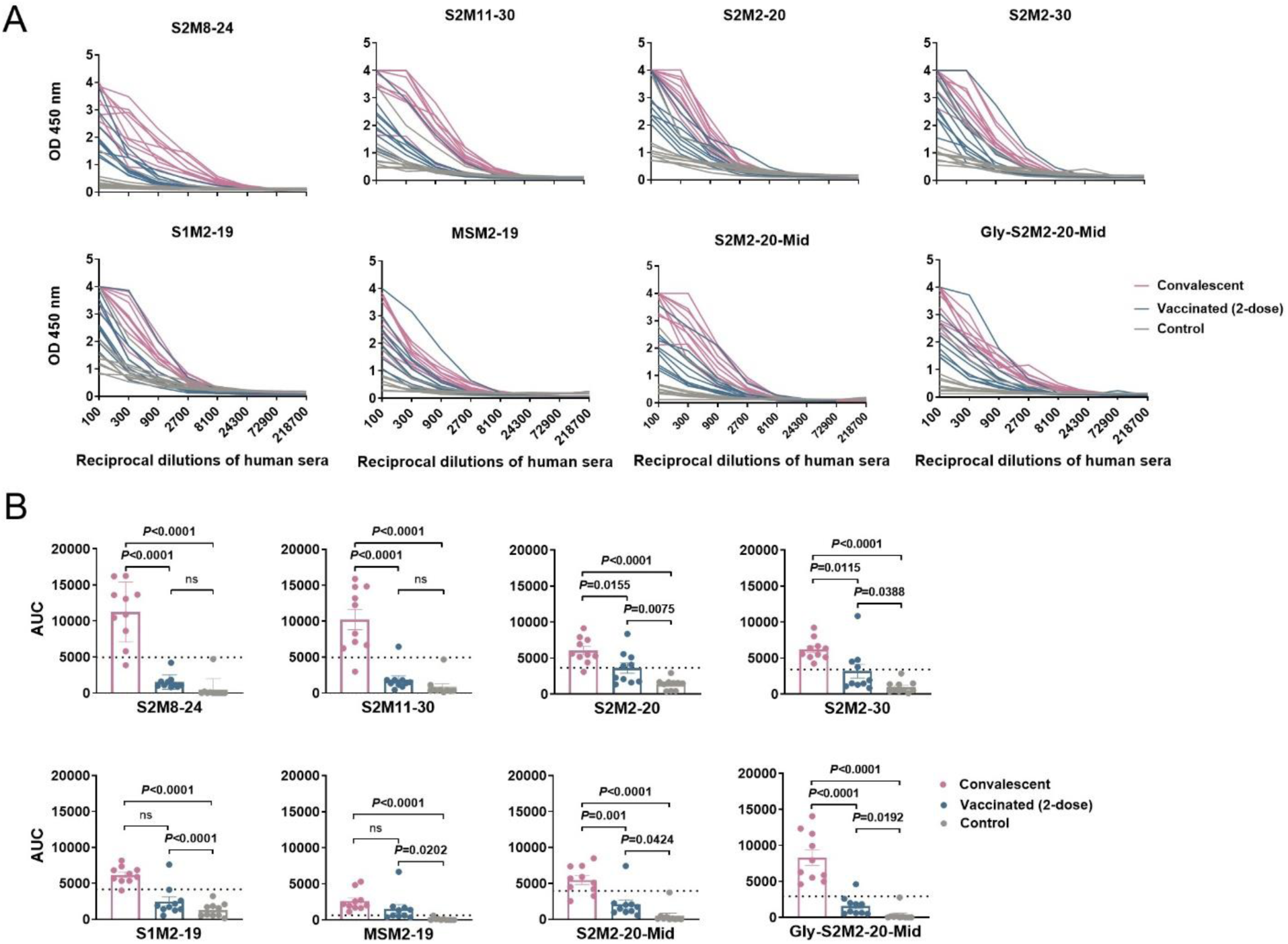
Serological evidence of M protein ectodomain-specific IgG responses in humans. (A) The human serum samples were collected from convalescent patients, vaccinees who ever received two doses of inactivated SARS-CoV-2 vaccines, and child outpatients in the middle of the year 2019 as negative controls. (B) AUC method interpretation of the same data as in (A). The dashed lines in the figures represent the Mean + 2 × SEM of the values from negative controls, and values above the dashed lines are regarded as positive results. Statistical significance between compared groups was determined using an unpaired t-test. P values less than 0.05 are considered statistically significant and ‘ns’ means not significant.

**Figure S2.**
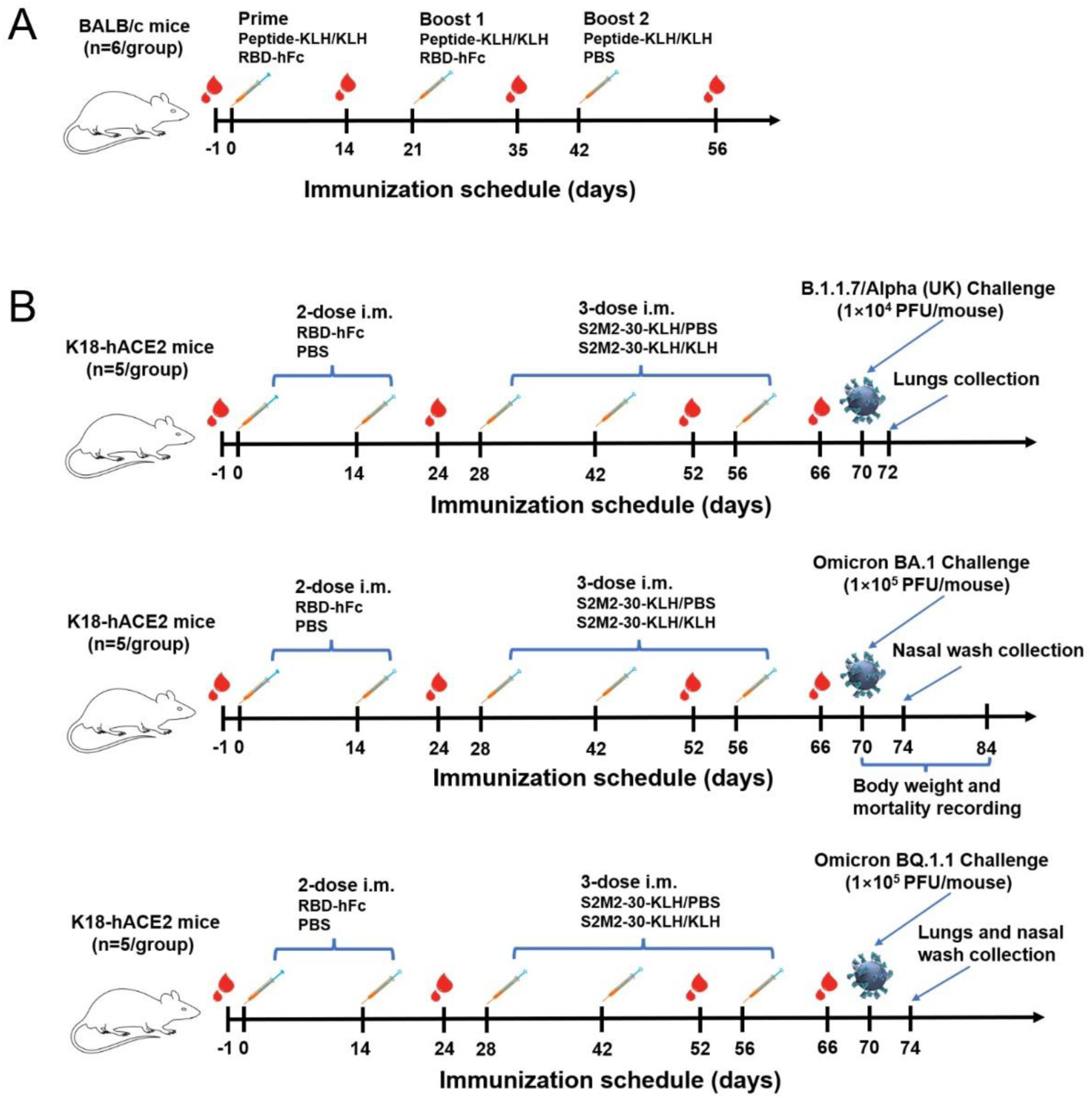
The regimens of immunizations and viral challenges.

**Figure S3.**
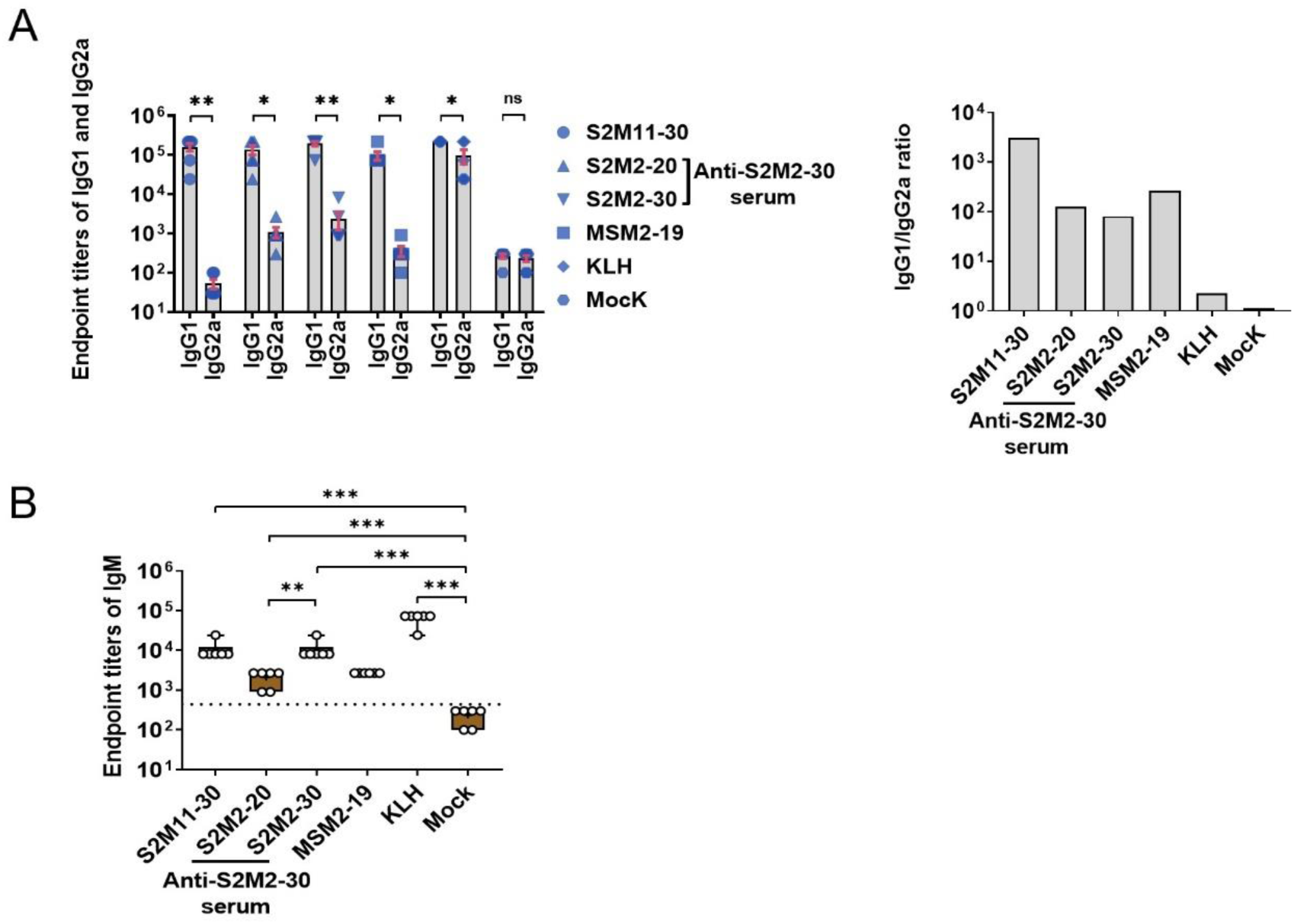
Endpoint titers of peptide-specific IgG1, IgG2a, and IgM. (A) Titrations of peptide-specific IgG1 and IgG2a. (B) Titration of peptide-specific IgM. The S2M2-30-KLH immune serum was titrated against both S2M2-20 and S2M2-30 peptides in ELISA. Data represent Mean ± SEM. The dashed lines in the figures represent the Mean + 2 × SEM of the values from negative controls, and values above the dashed lines are regarded as positive results. Statistical significance was determined using an unpaired t-test. P values less than 0.05 are considered as statistically significant and the *, **, and *** represent P values less than 0.05, 0.01, and 0.001, respectively. The ‘ns’ indicates not significant.

**Figure S4.**
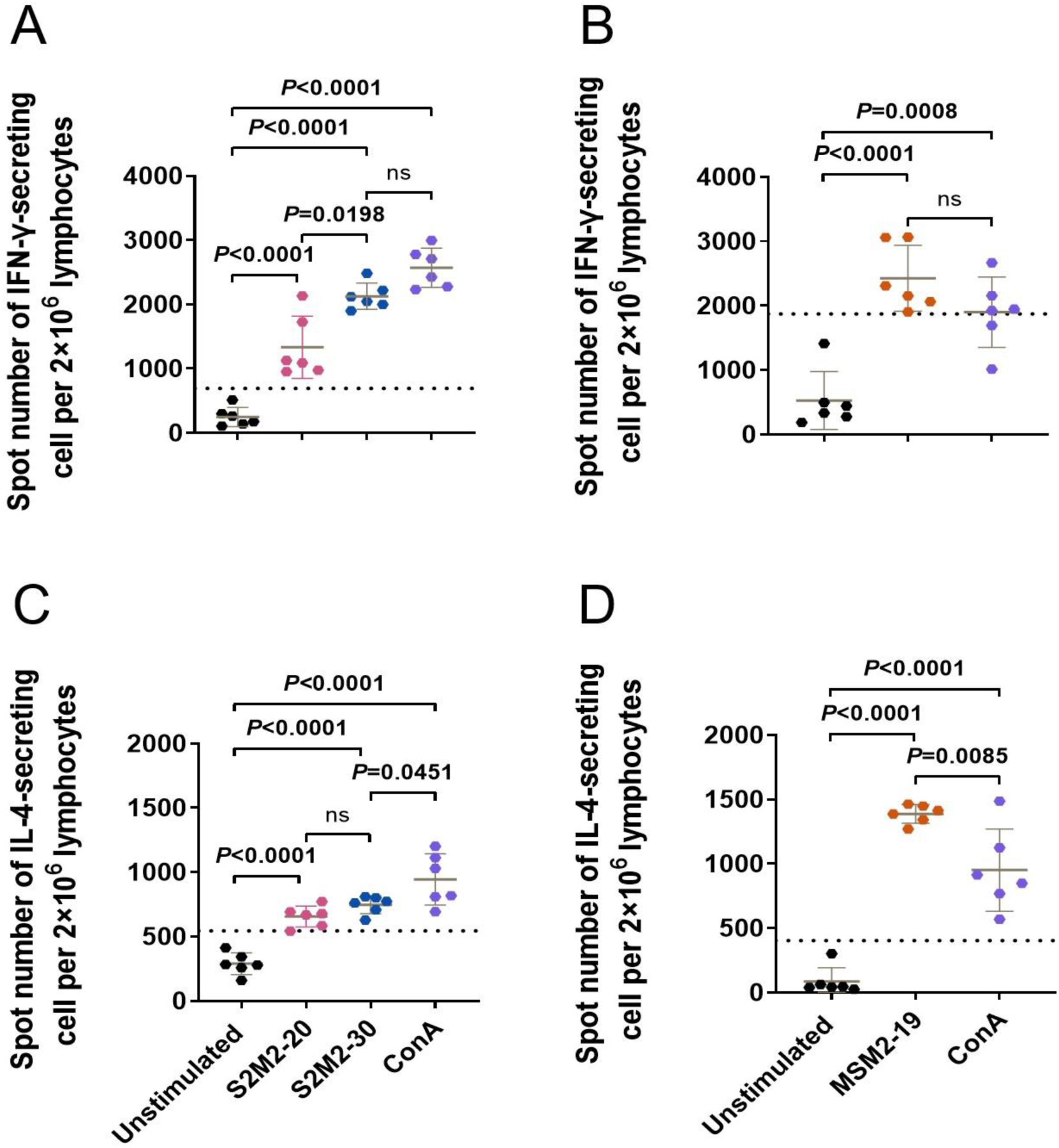
ELISPOT assays. Quantification of (A, B) IFN-γ-secreting and (C, D) IL-4-secreting splenocytes from the mice immunized with (A, C) S2M2-30-KLH and (B, D) MSM2-19-KLH immunogens, respectively. The unstimulated and ConA-stimulated cell samples were used as negative and positive controls, respectively. Data represent Mean ± SEM. The dashed lines in the figures represent the Mean + 2 × SEM of the values from negative controls, and values above the dashed lines are regarded as positive results. Statistical significance was determined using an unpaired t-test. P values less than 0.05 are considered statistically significant and the ‘ns’ indicates not significant.

**Table S1.**
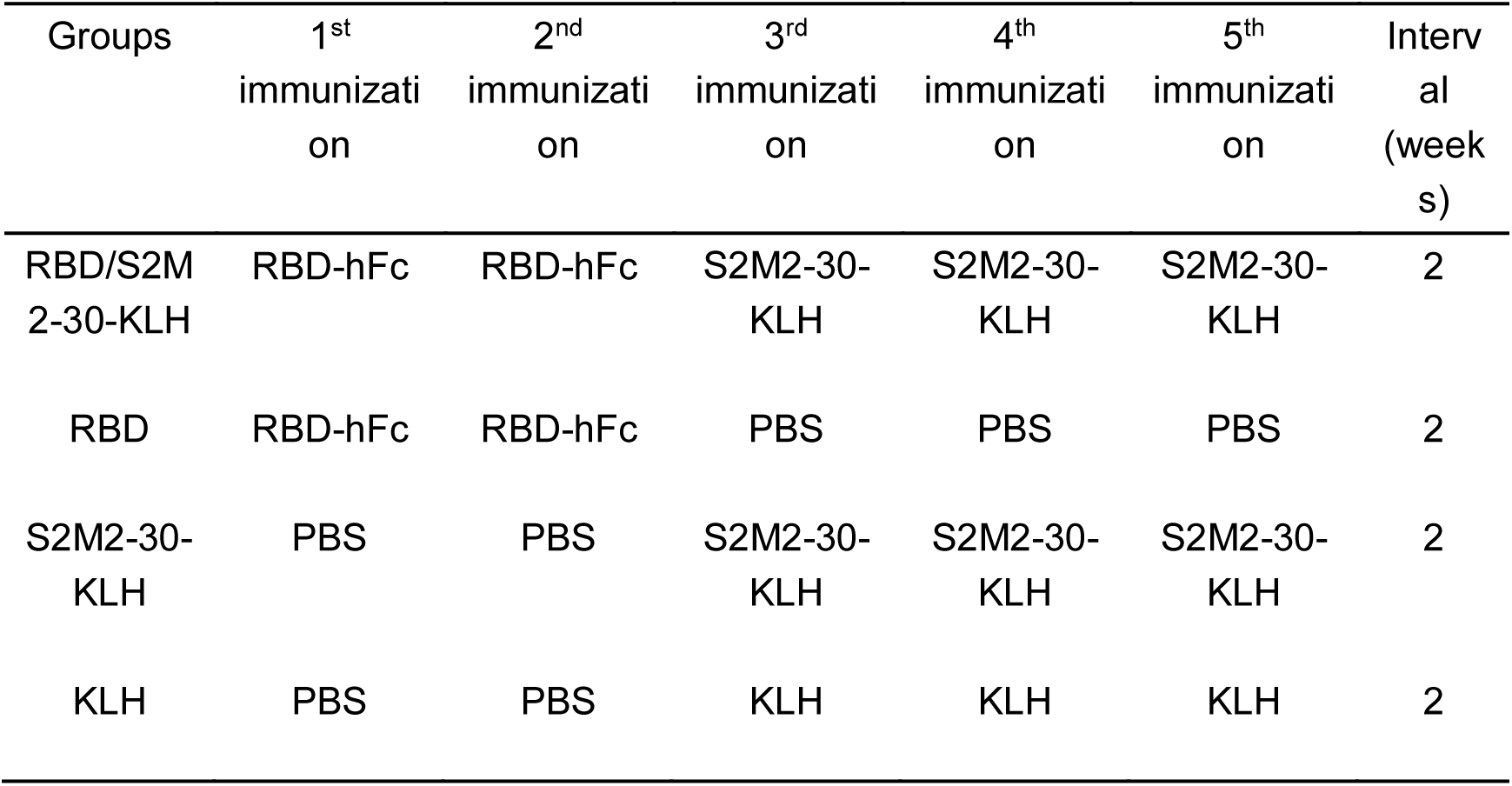
K18-hACE2 mice immunization regimens.

| Groups | 1<sup>st</sup><br>immunizati<br>on | 2<sup>nd</sup><br>immunizati<br>on | 3<sup>rd</sup><br>immunizati<br>on | 4<sup>th</sup><br>immunizati<br>on | 5<sup>th</sup><br>immunizati<br>on | Interv<br>al<br>(week<br>s) |
| --- | --- | --- | --- | --- | --- | --- |
| RBD/S2M<br>2-30-KLH | RBD-hFc | RBD-hFc | S2M2-30-KLH | S2M2-30-KLH | S2M2-30-KLH | 2 |
| RBD | RBD-hFc | RBD-hFc | PBS | PBS | PBS | 2 |
| S2M2-30-KLH | PBS | PBS | S2M2-30-KLH | S2M2-30-KLH | S2M2-30-KLH | 2 |
| KLH | PBS | PBS | KLH | KLH | KLH | 2 |

